# Two Distinct Attentional Priorities Guide Exploratory and Exploitative Gaze in Parallel

**DOI:** 10.1101/2025.09.20.677503

**Authors:** Xuan Wen, Alireza Seyed Hassani, Adam Neumann, Paul Tiesinga, Thilo Womelsdorf

## Abstract

Gaze is directed to visual objects that are informative, reward-predictive, or novel. These gaze preferences may reflect the parallel influence of two separable attention systems: Exploratory attention prioritizing uncertainty and exploitative attention prioritizing learned information about reward. We tested this hypothesis in nonhuman primates learning feature-based attention to objects that had either previously learned reward associations or were novel. The reward history of features slowed down learning by attracting fixations of non-rewarded distractors that were previously targets. This reward history bias persisted in fixations used to choose objects even after choice accuracy stabilized. In contrast, fixational sampling that preceded a choice showed negligible history biases that were overcome quickly in favor of wider exploratory sampling. Quantifying the exploratory value object features with a Parallel Belief States model of attention confirmed that exploratory fixational sampling was unaffected by reward history, while exploitative fixations that committed to a decision showed persistent target history biases. These findings suggest that gaze is guided by two separable attentional priorities in parallel. Exploratory attention prioritizes uncertain items and instantiates information sampling, while exploitative attentional priority guides gaze to current and previously goal-relevant features.

## Introduction

Gaze is directed to objects that have predictive reward value (Ghazizadeh et al., 2016; De Tommaso et al., 2019; Le Pelley et al., 2019), are informative even when not linked to rewards (Ghazizadeh et al., 2016; Ludwig and Evens, 2017; Pearson et al., 2024; Chow et al., 2025), or are novel (Monosov, 2024). These different types of gaze suggests that gaze is used to balance opposing functional needs to explore cues whose reward value is uncertain, while also exploiting knowledge about which cues are maximally predicting reward (Beesley et al., 2015; Easdale et al., 2019). The distinction of gaze to reflect exploration versus exploitation raises the question how separable the representations are that guide eye movements to satisfy the functional needs of exploring and exploiting (Tatler et al., 2011). Here, we hypothesized that exploratory and exploitative eye movement originate from separable systems that act in parallel to explore and exploit individual visual features during the deployment of feature-based attention.

A long-standing perspective suggests that visual attention predominantly guides eye movements towards uncertain reward associations, reflecting exploratory sampling of visual information (Pearce and Hall, 1980; Chow et al., 2025). According to this framework, fixational sampling of objects is guided by the uncertainty about the value of objects either because they are novel or resulted in unexpected outcomes in a recent encounter. Once the value of objects become predictable, this *exploratory attention* system is substituted by a separate system that guides fixations and choices towards stimuli that are maximally reward predictive (Gottlieb, 2012). The reward-driven system can be considered to reflect *exploitative attention* because it is driven by reward expectations (Easdale et al., 2019), and also by subjective choice preferences that are more typically studied in economic decision-making tasks (Le Pelley et al., 2019; Yang and Krajbich, 2023). The distinction of exploratory versus exploitative attention implies that separable types of attentional priorities guide eye movements towards objects. One priority map evaluates the uncertainty of objects and favors objects that are informative, while another priority map favor objects that have high values for the behavioral goal of a subject. It has remained unclear, however, whether these two types of attentional priorities reflect the same system that transitions from an exploratory mode to an exploitative mode once reward values become more predictable, or whether they are two segregated systems that act in parallel. Here, we devised a multidimensional feature-based attention learning task that suggests these two systems act separable, but act in parallel act in parallel on different individual visual features so that some features are attended based on their reward predictability, while other features are attended because their values are uncertain.

Characterizing exploratory and exploitative attention requires determining how they relate to the goal-based and the experience-dependent attention systems that previous work established in addition to the bottom-up saliency based attention system (Awh et al., 2012). Goal-based attention is reflected in a priority map of goal-relevant features that guides attention towards stimuli with features matching the prioritized target (Womelsdorf and Everling, 2015; Bisley and Mirpour, 2019). Such goal-based attention will correspond to exploitative attention towards features that are known targets, and is distinct from exploratory attention which is driven by not knowing the relevant features. The second experience-dependent attention systems explains the pervasive effects of selection history and refers to the systematic bias of directing attention to stimuli that were previously rewarded or chosen even if these stimuli have become non-rewarded distractors (Awh et al., 2012; Anderson et al., 2021). Selection history biases typically impair visual search of visual features that were previously defining a target object but have become distractors (Kyllingsbaek et al., 2001; Kyllingsbaek et al., 2014; Miranda and Palmer, 2014; Lin et al., 2016; Sha and Jiang, 2016; Qu et al., 2017). Selection history effects persist over hundreds of trials in visual search studies, reflecting a habitual, involuntary selection of non-rewarded distractor objects that neither satisfy an exploratory nor an apparent exploitative purpose, although they appear to signify exploiting previously learned stimulus-response mappings (Anderson et al., 2021).

Based on these considerations the goal-based attention system is well separable from an exploratory attention system, and the experience-dependent (*selection-history based*) attention systems might constitute a third system that biases attention independently of the other two systems. Here, we tested whether the influence of experience-dependent attention can be conceived of as independent of the other two systems, in which case selection history effects should affect both, exploratory and exploitative attentional orienting. Contrary to this suggestion we found that the exploratory attention system is largely unaffected by selection history biases, suggesting that attentional priorities that guide exploratory attention are distinct from both goal-based and experience-dependent priorities. We arrive at this conclusion with a task that required subjects to either learn the reward association with a novel target feature or to update a target feature by re-assigning the reward associations from a feature that was previously a distractor feature. We found that updating target features is consistently more difficult than learning new targets and induces persistent biases towards previous targets that have become distractors. In contrast, exploratory eye movements to objects prior to choosing an object were largely unaffected by target history biases although they continued sampling objects even after subjects had fully learned which objects were new targets and distractors. These findings document a dissociation of attentional representations guiding exploratory versus exploitative eye movement during and after learning of feature-based visual attention.

## Results

We collected performance of two monkeys with a feature-based Learning and Updating task in 41 experimental sessions (subject I: 27, W: 14). The task required subjects to learn through trial-and-error which object feature is the rewarded target feature in blocks of 30-50 trials. Each trial presented three objects that differed in features of two feature dimensions (e.g. different colors and different shapes), i.e. there were six distinct features in each trial, one of which was the target feature (**Fig. 1A**). Subjects chose an object by fixating it for 0.7 s, which led to visual and auditory feedback and to earning visual tokens if the object contained the target feature, or the loss of token(s) if the chosen object did not contain the target. Visual tokens were added or subtracted from a token bar on top of the screen and cashed out for fluid reward once eight tokens were collected. After initiating a trial, subjects had up to 5 s to freely look at objects prior to committing choosing an object. We defined one feature as the correct target feature and five other features as distractor features for an entire block. When a block switched, object features were either novel (e.g. different sets of colors and shapes), or the same features of the previous block were used but a different feature was assigned the rewarded target feature (*Novel* vs. *Same* block conditions, **Suppl. Fig. S1**). Blocks also differed in whether the new target feature was from the same or different feature dimension as the previous blocks target, whether subjects received 2 or 4 tokens for correct choices, and whether they lost 1 and 3 token for incorrect choices (**Fig. 1B**).

**Fig. 1.**
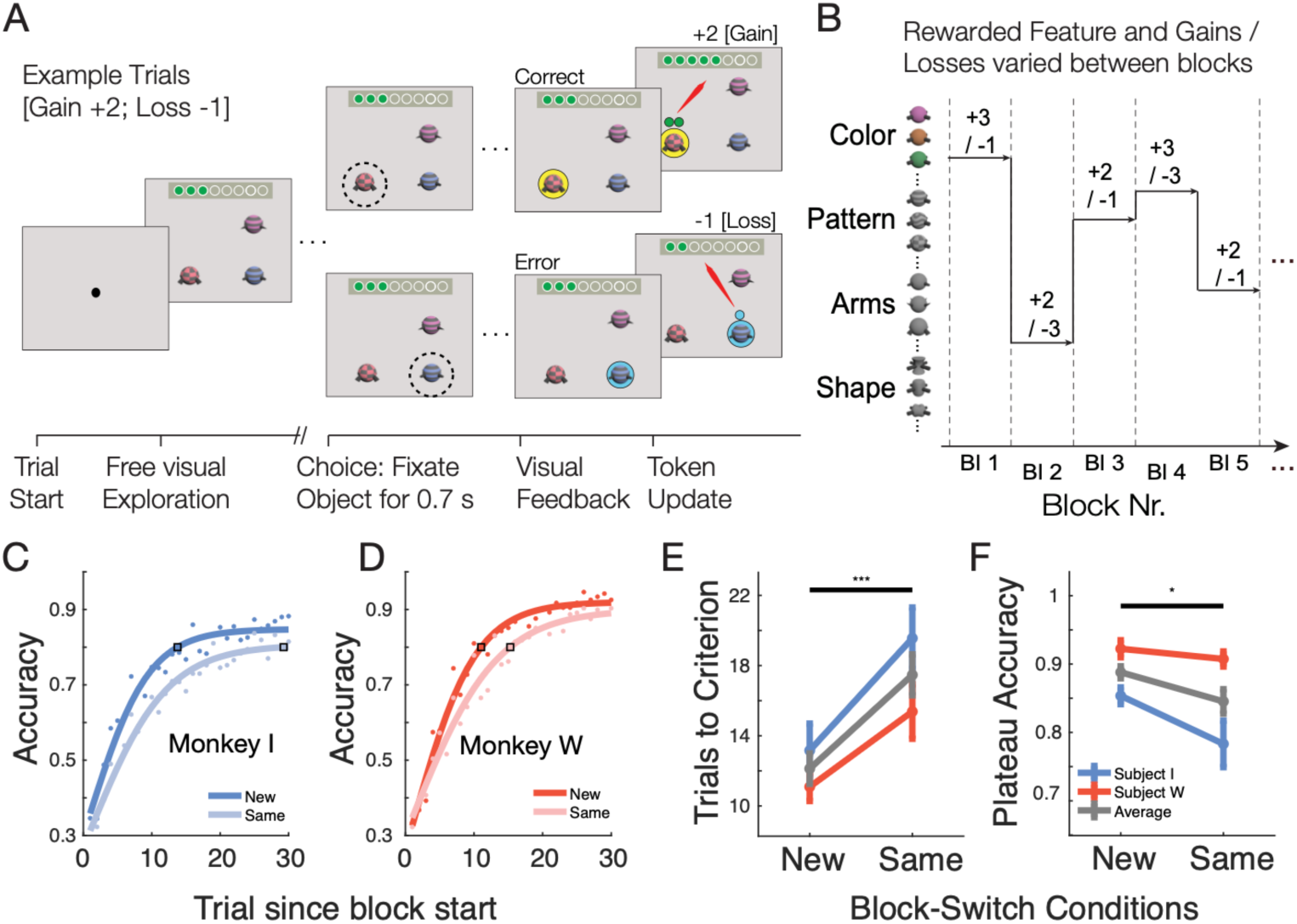
Feature-based learning & updating task paradigm. (**A**) Each trial allowed <5 s for visually exploring three objects. Subjects chose an object by fixating it for 0.7 s, which was followed by a visual halo around the chosen object and the tokens gained for correct or lost for incorrect choices. (**B**) Within blocks of 30-50 trials one feature was the rewarded target, and the motivational context was stable (e.g. +3 tokens for correct, and −1 token for incorrect choices). A new block had either new features or used the same features as the previous trial with reassigned target feature (*see* **Suppl. Fig. S1**). (**C,D**) Proportion of correct choices for subject I (*C*) and W (*D*) increased over trials slower when the new target feature was a distracting feature in the previous trial (*Same*) as opposed to a new feature (*New*). Open square demonstrates the trial at which 90% of maximal plateau performance was reached which was the criterion defining learning. (**E**) Learning was slower in the *Same* than *New* block transitions. (**F**) Plateau performance was lower in the *Same* than *New* block transitions. Stars denote significance level.

### Target and distractor history slows learning of feature-based priority

Subject completed on average 34.85 blocks per session and 37.43 trials per block (I: 925 blocks, 38.56 trials per block; W: 504 blocks, 35.35 trials per block). Across all conditions, accuracy started at chance level and reached 80% on average within 14.82 trials (I: 17.18 ± 1.21; W: 13.55 ± 1.15), with a plateau of 86.7% accuracy after 30 trials (I: 86 ± 3; W: 91.3 ± 1). Both subjects reached the learning criterion of 80% later when the block showed the same features as the previous block as opposed to novel object features (**Fig. 1C-D**), slowing down learning by on average 5.343 trials (Same: I: 19.56 ± 1.75; W: 15.38 ± 1.55; New: I: 13.14 ± 1.53 ; W: 11.09 ± 0.81) (**Fig. 1E**). Subjects also reached lower plateau performance with the *Same* as opposed to *Novel* features, suggesting that there was a persistent influence of the history of target- and distractor-features from the previous blocks (Same: I: 78% ± 3; W: 91% ± 1; New: I: 85% ± 1; W: 92% ± 1) (**Fig. 1F**). The slower learning and poorer performance was specific to the *Same* versus *Novel* block condition comparison and not secondary to extra- vs intra-dimensional target changes (**Suppl. Fig. S2A-F**) or to motivational factors. Blocks with higher vs lower motivational incentives (4 vs 2 tokens gained for correct responses) or lower and higher symbolic punishments (−1 vs −3 tokens lost for incorrect responses) did not systematically affect learning speed and plateau accuracies (**Suppl. Fig. S2E-L**).

### Target history effects are persistent in choice but transitory in fixational sampling

Poorer learning in the *Same* versus *Novel* block condition suggests that the stimulus history slowed down learning. To quantify the effect of stimulus history we compared the relative proportions of errors made by choosing distractor objects with novel features in the *Novel* block condition, with choices of distractor objects having features that were rewarded targets in the previous block (*previous target features*), and with choices of objects with features that were already distractor features in the previous block (*previous distractor features*). Choices of distractors with features that were previous targets were more prevalent than choices of previous distractor or novel features (New vs. Prev. Target: t = −10.41, p < 0.001; Prev. Target vs. Prev. Distractor: t = 9.15, p < 0.001), while erroneous choices of objects with novel features were more likely than of previous distractor features (Prev. Target vs. Prev. Distractor: t = 14.77, p < 0.001) (**Fig. 2A**). These target-history and distractor-history effects were persistent across all trials of the learning block (**Fig. 2C**). Choosing the previous target remained at a similar level compared to other distractors throughout the block (linear regression slope r = 0.31, n.s.) while erroneous choices of previous distractors continued decreasing with trials in a block (linear regression slope r= −0.45, p = 0.011).

**Fig. 2.**
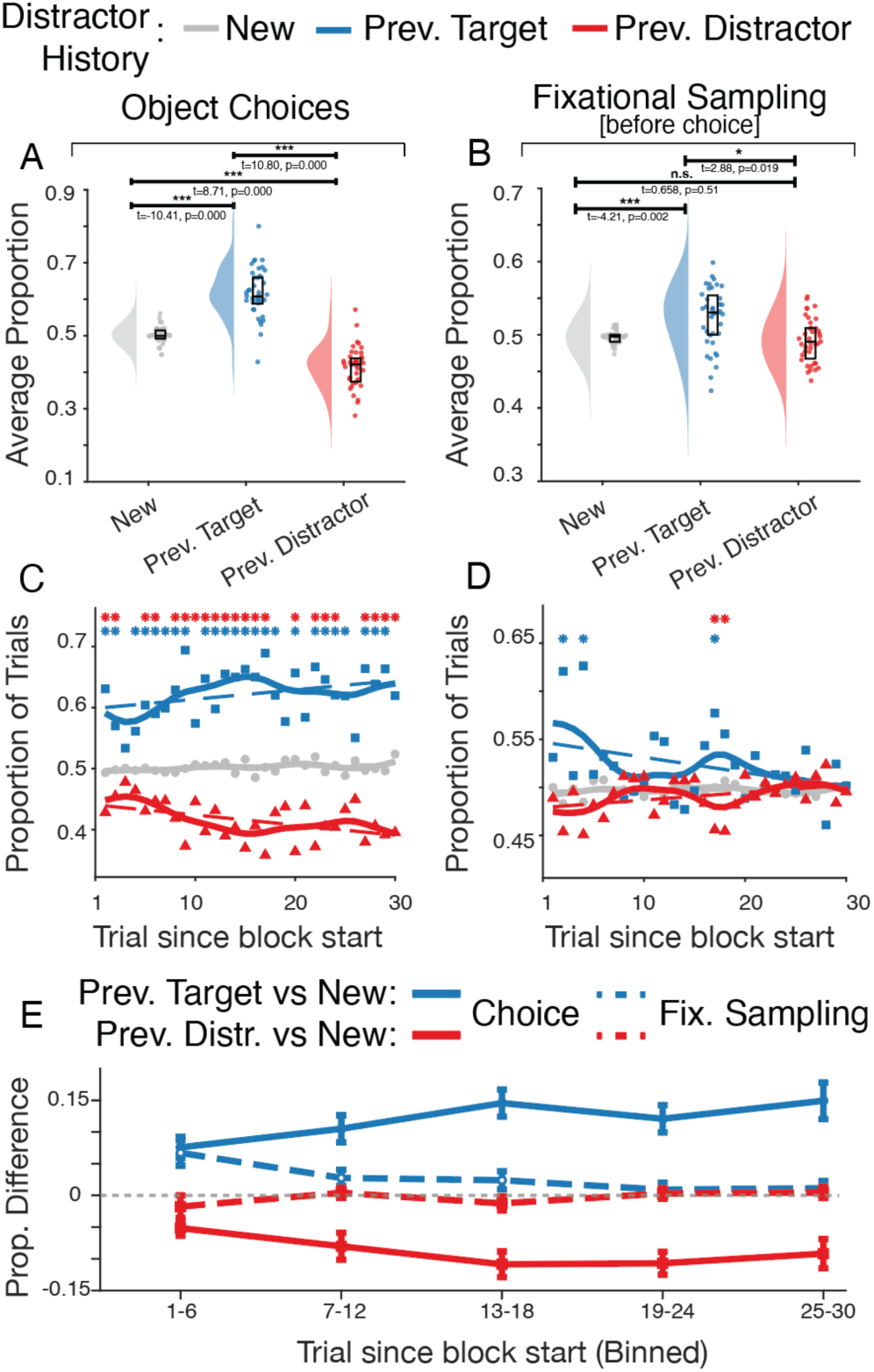
Selection-history effects on choices and fixational sampling. (**A**) Average proportion of trials (±SE) in which subjects erroneously chose a distractor object with features that were *New* (that were not used in the previous block), or were Previous Target features or Previous Distractor features in the last bock. Stars denote a two-sample t-test with a Bonferroni correction (**B**) Same as *A* but only counting fixations on objects that occurred prior to the last fixation of the chosen objects. (**C**,**D**) The same as *A* and *B* but separately quantified for individual trials in a block since block start. Solid lines are cubic spline interpolated data. Dashed lines are linear regressions. Stars at the top denote statistical difference of the previous target (red) and previous distractor (yellow) data points relative to new distractor using two-proportion z-test. (**E**) The difference of the proportion of choices (±SE) for previous target (red) and distractor (blue) features relative to new features binned in consecutive, nonoverlapping sets of five trials.

Target- and distractor- history effects are considered a major source of attentional top-down control, but the pervasiveness of selection-history effects on attention has remained unclear and is under-investigated (Anderson et al., 2021). If selection-history has a pervasive influence on controlling which object is attended, it should be evident in overt attentional sampling of objects with eye movements irrespective of the act of choosing an object. We tested this assumption by analyzing how often subjects looked at distractor objects prior to making a choice. Overall, the fixational sampling of distractors showed the same target- and distractor-history biases as the actual choices of objects. Fixational sampling of distractors with features that were previous targets were more prevalent than choices of previous distractor features (Prev. Target vs. Prev. Distractor: t = 3.86, p = 0.001), while fixational sampling of objects with novel features was more likely than of previous distractors (New vs. Prev. Target: t = −4.14, p < 0.001) (**Fig. 2B**). However, the time course of these effects differed to the time course of selection history effects in actual choices. Selection history effects in fixational sampling were less persistent and largely limited to a few trials at the beginning of a block, which was evident in a negative slope of a regression of the effect over trials (Prev. Target: linear regression: r = −0.37, p = 0.04)(**Fig. 2D**). We quantified the difference of fixational sampling by averaging the difference of the proportion of trials with fixational sampling of previous targets to novel objects in consecutive six trials across the block and found that fixational sampling of previous targets were more prevalent than novel objects during the first twelve trials and not detectable thereafter (trials 1-6: difference = 0.067±0.020, t = 3.34, p = 0.0019; trials 7-12: difference = 0.028±0.013, t = 2.18, p = 0.035; trials 13-18: t = −1.67, p = 0.102) (**Fig. 2E**). There were no differences of fixational sampling of objects that were previous distractors versus new objects (**Fig. 2E**).

### Prolonged fixational sampling preceding erroneous choices

The previous finding raises the possibility that compared to choosing an object, fixational sampling is less persistently influenced by selection history and more quickly adjusts to explore objects when reward contingencies change during a block switch. To address this possibility, we analyzed the features of objects subjects fixated during a trial prior to the final fixation of the object they chose on that trial (**Fig. 3A**). Subjects may start a trial and then directly look at and choose the object with target or a distractor features (*Tc* or *Dc*, ‘c’ marking the choice fixation), or they may fixate other objects with distractor (*D*) or target features (*T*) prior to choosing an object, such as looking at a distractor before looking at and choosing the target (sequence *D-Tc*) or at the target before choosing a distractor (*T-Dc*) (**Fig. 3B**, **Suppl. Fig S3A**). In the following we focus on the last 2 trial fixations because trials with >2 fixations prior to a choice (e.g. D-T-Dc, or D-D-Dc) were rare and overall similar to trials with two fixations (**Suppl. Fig 3B,C**).

**Fig. 3.**
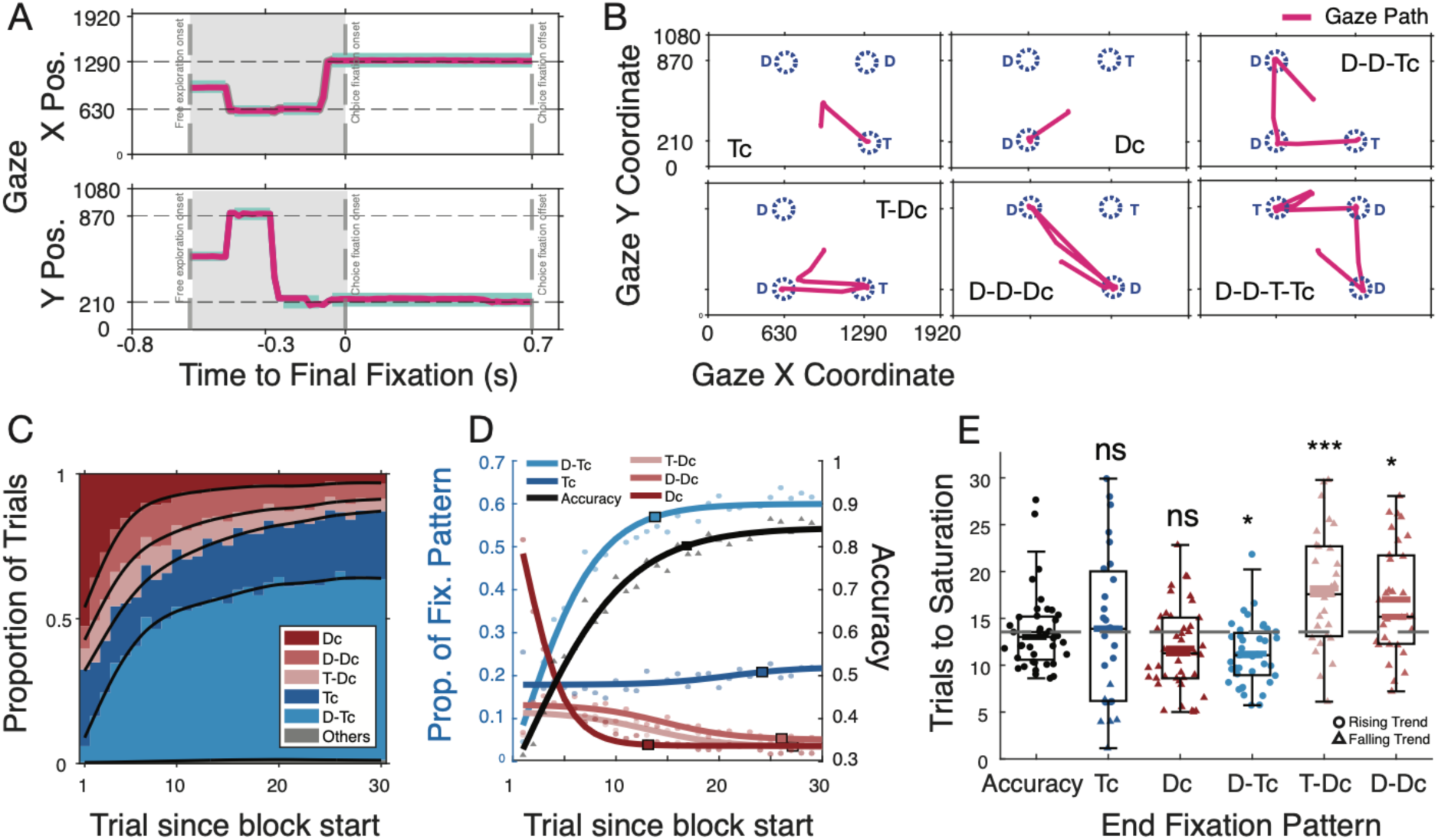
Fixational Sampling sequences and comparison to accuracy. (**A**) Example eye position traces for X and Y positions (upper and lower) in a trial relative to the onset of the choice-fixation that was maintained for 0.7 s. The grey shades area marks the free visual exploration trial and shows a saccade towards the upper right object, then a downwards saccade to the lower left object position, and then a saccade to the left where the fixation stayed for >0.7s (**B**) 2-Dimensional fixational sampling sequences in six example trials. Fixations start in the center (which initiated the trial) and subjects could made any fixational sequences in <5 s before choosing an object by sustaining fixation for 0.7 s. (**C**) Proportion of trials for different fixation sequences considering trials with 2 or less fixations. Tc and Dc denote choice fixations, T and D denote fixational sampling prior to a choice. See **Suppl. Fig. S3** for sequences with >2 fixations. (**D**) Proportion of fixational sampling sequences (left y-axis) and choice accuracy (right y-axis) across trials in a block. Black squares denote trials at which half sigmoid reached 90% to saturation. (**E**) Trials-to-reach plateau performance (learning speed) for accuracy (*left*) and different fixational sequences (*right*) across sessions. Stars denote significance level using a two-sample t-test.

Comparing the fixational sampling patterns over trials in a block with the change of choice accuracy revealed various notable differences (**Fig. 3C,D**). We found that while choice accuracy plateaued at trial 13.55 (±0.65), the proportion of trials in which subjects fixated and immediately chose the target object (*Tc*) without additional fixations remained rather constant across trials suggesting that the learning process is reflected in changes of the fixational sequences rather than in an increase of fast *Tc* choices (**Fig. 3D**). Of those fixation sequences, the *D-Tc* sequence, that indexes fixating and then rejecting a distractor to then choose the new target, increased most rapidly, plateauing significantly earlier than choice accuracy at trial 11.23 (±0.54; t=2.53,p=0.013; **Fig. 3E**). In contrast, fixational sampling of a target or of distractors that ended with choosing a distractor object (*T-Dc* and *D-Dc* sequences) continued to decline at a slower time course than choice accuracy (**Fig. 3D,E**). The plateau for the *D-Dc* sequence occurred at trial 17.00±1.05, and for the *T-Dc* condition, it occurred at trial 18.21±1.16, which both were later than the 13.55 trials needed to reach the plateau for choice accuracy (*D-Dc* vs choice accuracy: t=−2.59, p=0.012; *T-Dc* vs choice accuracy: t=−3.88,p<0.001) (**Fig. 3E**).

### Fixational sampling reflects feature-specific uncertainty

The prolonged fixational sampling of objects on error trials (*D-Dc* and *T-Dc*) (**Fig 3E**) may indicate continued uncertainty about the relevance of objects, but these additional eye movements on error trials may also reflect the influence of target-history on eye movements. To disentangle the effects of target history and uncertainty on fixational sampling we estimated the uncertainty of features in each trial with a model that tracks the certainty of subjective beliefs about the certainty about target features. The model was introduced as a Parallel Belief States (PBS) model that assumes subjects tracks how relevant visual features are for attention and choices by updating the subjective certainty, or belief, about their relevance (Goudar et al., 2024; Tiesinga and Womelsdorf, 2025). The belief is quantified as a hidden state variable in a Hidden Markov Model that estimates the likelihood that a feature will be chosen given the recent history of rewards and choices (**Fig 4A**). We applied the PBS model to choices of subjects and found that they were accounted for by assuming each object feature can be in one of three separable belief states that signify whether the feature is either (1) avoided with high certainty that it is a distractor and unrewarded (Avoid State, *Avd*), (2) exploited with a high certainty of being rewarded and chosen (Exploit State, *Ept*), or else (3) explored and being considered to be chosen because of uncertainty about its reward value (Exploration State, *Exp*) (*see* Methods). Consistent with these states, features of distractor objects were more likely in Avoid and Exploration states whereas objects with the target feature had more likely features in the Exploit state (**Fig. 4B**). Beyond the label of targets and distractors, the model states closely predicted whether an object will be chosen and fixated: Objects were chosen most likely when they had features in an Exploit state and less when both features were in Exploration states (With Ept: 0.9842±0.0067 vs. Without Ept: 0.1587±0.0624; p<0.001) (**Fig. 4C**). A key prediction of the model is that the Exploration state of features reflects uncertainty about the relevance of a feature. We hypothesized that if uncertainty drives attention, than there should be more fixational sampling to objects with features in an Exploration state. We confirmed this prediction. Objects were fixated (prior to choosing an object) most likely when one or both features were in an Explore state and less when they were in an Exploit or Avoid state (With Ept: 0.1829±0.0451 vs. Without Ept: 0.4614±0.0327; p<0.001) (**Fig. 4D**).

**Fig. 4.**
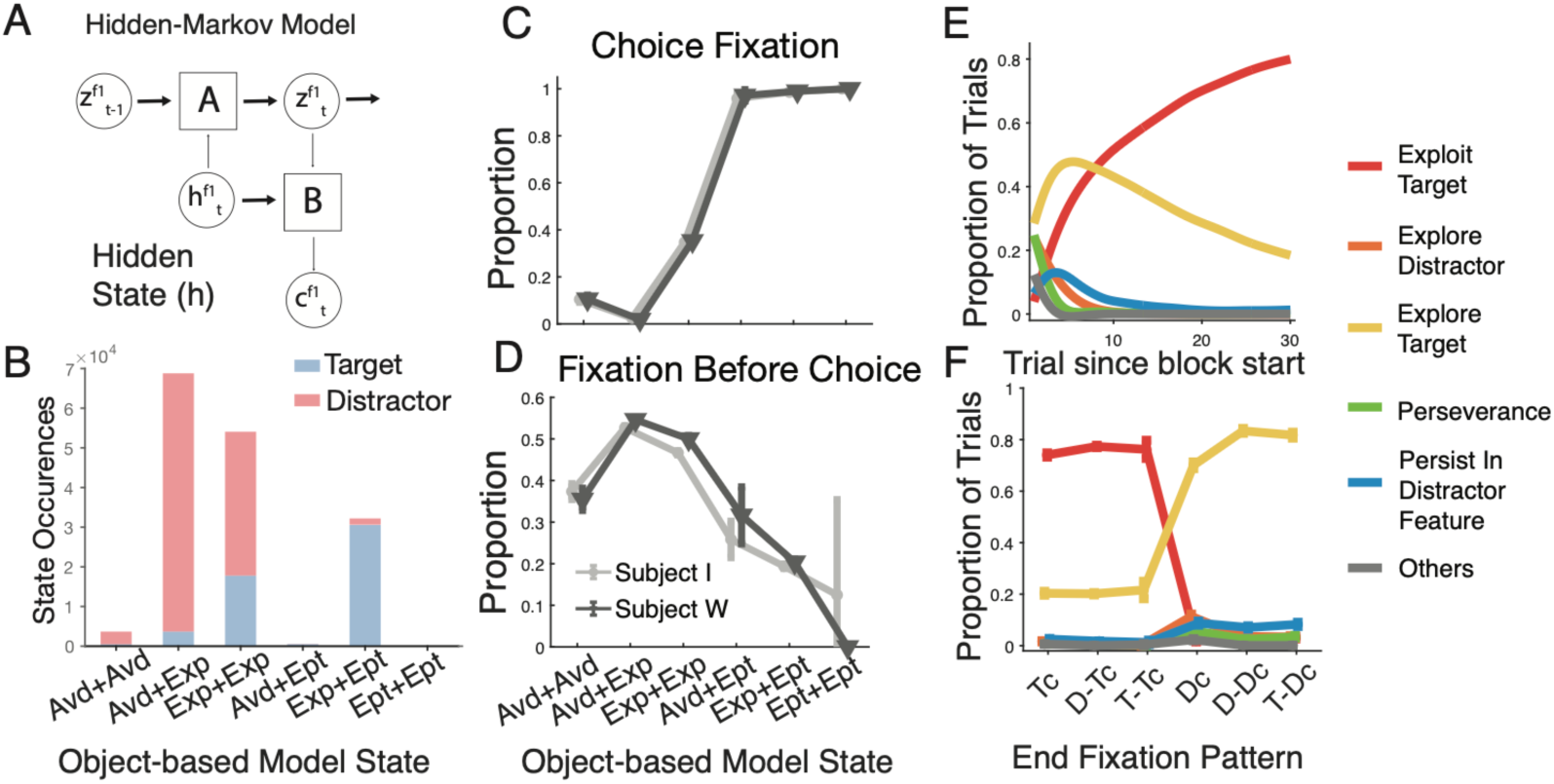
Feature-specific belief states underlying choices and fixational sampling characterize behavioral learning progress. (**A**) A hidden Markov model estimates how likely an object feature is being chosen and rewarded using hidden state variable h and previous trials’ choices and rewards (*see* Methods). The likelihood estimate corresponds to discrete belief states for each feature to be either irrelevant (Avoid state), relevant (Exploitation state) or with uncertain relevance (Exploration state). (**B**) State occurrences of features on target objects (*blue*) and distractor objects (*red*) (*x-axis*). As expected, target objects had more likely a feature in the Exploit and Explore state. (**C**,**D**) The proportion of object states for objects that were chosen (*C*) or fixated prior to a choice (*D*) for each monkey (grey shading). Fixational sampling was more likely when an object feature was in Exploration state. (**E**) Classification of trials into six different states according to the state of the target feature in the current and previous trial (*see* text). Trials in ‘Explore Distractor state’ declines rapidly early in a block, while the state ‘Persist in distractor features’ (blue) more gradually declined. ‘Explore target states initially increased and then leveled off as the ‘Exploit target state became more prevalent. (**F**) Proportion of trial state occurences calculated separastley for each end-fixational sampling pattern (*x-axis*) shows that distractor objects are sampled mostly when trials are in an ‘Explore State, but also in trial states ‘Persist in distractor feature’.

### Tracking feature-specific beliefs recovers the learning progress

The previous analysis validates that the belief states of the PBS model account for choices when features were in an Exploit state, and for fixational sampling when features were in an Exploration state. We next tested how the parallel feature states change with learning and how target history affects these states. Each trial presented three objects that varied in six separate features. Considering that each feature could be in one of three distinct belief states this spans 3^6^ possible state combinations. To simplify the state space and make it interpretable we classified each trial according to the state of the rewarded target feature of the current and previous trial (**Fig. 4E**, *see* Methods). We found that considering five states provides a good overview of the learning progress. Trials in which the target, but not other features are in the Exploration state (‘*Explore Target*’ trial state) increased for the first few trials and then decreased, while trials in which the target feature is in the Exploit state (’*Exploit Target*’ trial state) increased steadily over trials (**Fig. 4E**). Conversely, trials in which distractor features, but not the target feature, were in an Exploration state (‘Explore Distractor’ trial state) rapidly declined at the beginning of a block. The classification of behavioral states over trials also recovered a rare state in which distractor features were in an Exploit state and the target feature was in Avoid or Explore states. We labeled this trial state ‘Persist in Distractor Feature’, as it reflects that subjects erroneously exploit distractor features. It constituted up to 10.59% of the trials early in the block (trials 1-6) before decreasing to 1.19% of trials throughout the remainder of a block (trial 25 to 30) (**Fig. 4E**).

Next, we calculated how likely these states occurred for each fixational sampling pattern. As expected, fixational sampling of the target was most likely in the Exploit Target and Explore Target State (**Fig. 4F**). In contrast, the majority of fixational sampling prior to an erroneous choice of the distractor occurred when the target feature was explored (Explore Target) and only a small minority of fixational sampling occurred when the distractors were erroneously exploited (Persist in Distractor Feature) (**Fig. 4F**). This pattern of result suggests that fixational sampling is only weakly affected by strong but wrong beliefs that a non-rewarded distractor is relevant.

### Fixational sampling is dominated by uncertainty, while choices reflect target history effects

The previous result indicates that fixational sampling might be less affected by wrong-held biases than actual choices. We directly tested this by comparing the proportion of choices and fixational sampling for objects that had features in the Avoid, Explore, and Exploit State. With regard to both, choices and fixational sampling, a neutral prediction is that target-history biases should be reflected in a higher proportion of choices as well as of fixational sampling of distractor objects with features that were previous targets regardless of the belief state of subjects about their relevance. However, this is not what we found. For choices, we found that erroneous choices of previous targets were more likely than previous distractors specifically in the Explore State (prev. targets vs prev. distractors, t= 19.26, p < 0.001, new vs. prev. targets, t = −10.475, p < 0.001) (**Fig. 5A**). In contrast, fixational sampling of objects did not show such a target history effect in the Explore State (n.s.) and showed only marginal differences in the Avoid state (n.s.) (**Fig. 5B**). These results were confirmed in the trial-by-trial analysis, which showed that selection history effects are evident in choices of subjects in the Explore States persistently throughout the block (**Fig. 5C**), consistent with the major behavioral effects (**Fig. 2**). In contrast, fixational sampling in the Explore state was void of target history biases (**Fig. 5D**). Together these results document that target history effects are predominantly driven by a system that commits to a choice, while fixational sampling is dominated by belief states about the uncertainty of the relevance of features.

**Fig. 5.**
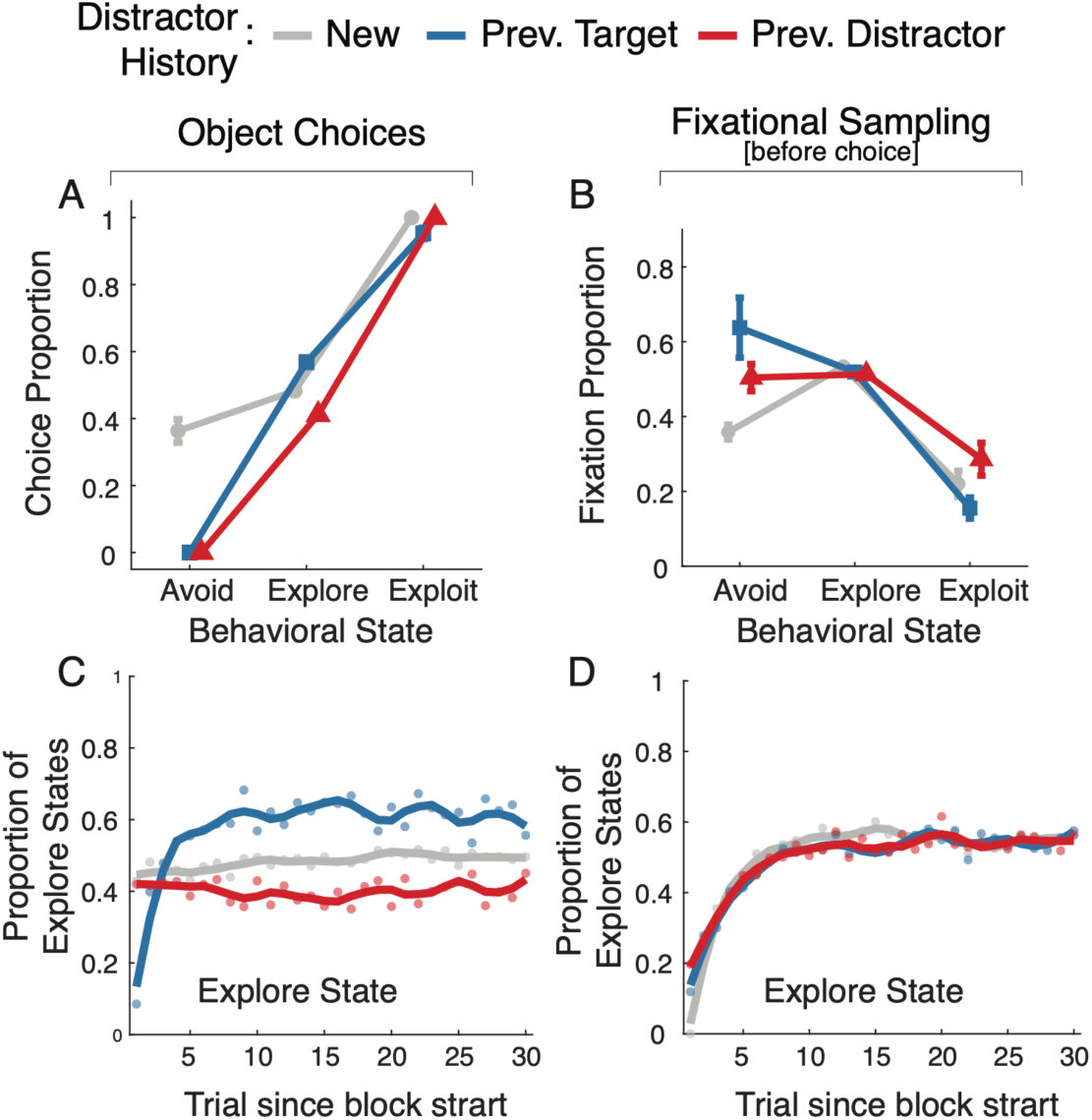
Fixational sampling during Exploratory Belief States are not affected by selection-history effects. (**A**) Proportion of choices of different distractor types in different behavioral states (x-axis). Previous targets are chosen more likely than prev. distractor objects in the Explore state (Mean ± 95%CI: Prev Target: 0.568 ± 0.012; Prev. Distractor: 0.4116 ± 0.009; two-sample t-test: t=19.26, Bonf. corrected p < 0.001). Please note that novel objects are more likely in the Avoid state at the beginning of the block because they lack choice and reward history, which likely accounts for them being similarly chosen in Avoid and Explore state. (**B**) Same format as A for fixational sampling. Here, previous targets are not differently chosen than previous distractor in the Explore state (Mean ± 95%CI: Prev Target: 0.519 ± 0.006; Prev. Distractor: 0.5133 ± 0.0052; two-sample t-test: t= 1.308, Bonf. corrected p > 0.1). (**C**,**D**) Same analysis as in *A+B* but for individual trials relative to the start of the block.

### Varying incentives and punishment avoidance do not interact with selection history biases

Target history effects might vary with the costs of choosing or sampling an unrewarded, distracting stimulus (Pearson and Le Pelley, 2020; Anderson et al., 2021). Choosing an unrewarded previous target is more costly when it results in loosing 3 than 1 token as it takes more time to compensate for the loss. We compared the target-and distractor-history biases between these *Loss* conditions and found that the average proportion of errors on previous targets was similar in both conditions (−1 token: 0.617±0.013 vs. −3 tokens: 0.611±0.014). Erroneous choices of previous distractors (−3 tokens: 0.423±0.012 vs. −1 token: 0.417±0.011, n.s.), and of novel distractors (−3 tokens: 0.495±0.006 vs. −1 token: 0.495±0.006, n.s.) were unchanged with high versus low symbolic punishment (**Suppl. Fig. S4A**). Fixational sampling prior to the choice also was unaffected by high versus low punishment for previous targets (−1 token: 0.521±0.010 vs. −3 tokens: 0.510±0.009), previous distractors (−1 token: 0.489±0.007 vs. −3 tokens: 0.499±0.007), or novel distractors (−1 token: 0.496±0.003 vs. −3 tokens: 0.493±0.003) (**Suppl. Fig. S4C**).

Next, we tested whether higher incentives to choose the correct target may reduce selection history bias, which would reflect that the magnitude of motivational incentives may strengthen attentional priority and weaken de-valued distractor representations. We compared previous target errors in conditions with high incentives for correct choices (3 versus 1 tokens gained) but found similar proportions of previous target choices (3 tokens: 0.629±0.014 vs. 2 tokens: 0.599±0.013, n.s.), previous distractors (3 tokens: 0.413±0.012 vs. 2 tokens: 0.426±0.011, n.s.), as well as novel distractors (3 tokens: 0.492±0.007 vs. 2 tokens: 0.498±0.005, n.s.) (**Suppl. Fig. S4B**). A similar null effect was observed in fixational sampling across previous targets (3 tokens: 0.519±0.009 vs. 2 tokens: 0.512±0.009), previous distractors (3 tokens: 0.489±0.007 vs. 2 tokens: 0.499±0.007), and novel distractors (3 tokens: 0.494±0.003 vs. 2 tokens: 0.495±0.003) (**Suppl. Fig. S4D**). The absence of a token gain effect is consistent with the conclusion of a recent meta-survey (Rusz et al., 2020) and reminiscent of studies reporting previous target effects on attention irrespective of the amount of monetary rewards in humans (Sha and Jiang, 2016; Grubb and Li, 2018).

## Discussion

We showed that gaze fixations of objects are differently affected by selection history biases depending on whether they sample objects prior to committing to a choice or whether they are used for choosing an object. Fixating to choose an object showed persistent biases towards non-rewarded distractors with features that were target features in previous blocks. In contrast fixational sampling of objects prior to choosing an object showed the same bias only weakly and transiently at the beginning of updating the rewarded target feature (**Fig. 2C-E**). This finding suggests that fixations that explore objects are governed by a representation that more quickly and flexibly overcomes reward history biases compared to fixation used to choose objects and exploit their predicted reward association.

The distinct influence of reward histories on exploratory and exploitative fixations became evident in a task paradigm in which animals were free to sample objects with brief fixations prior to committing to a choice. Behavior that is driven to optimize rewards would be expected to use the opportunity to freely sample objects during the early learning and updating period to establish feature-reward predictions for target and distractor features and, once learning completed, to quickly fixate and chose the most reward predictive object. However, this is not what was observed. Subjects sampled objects not only during the initial learning period, but continued to direct eye movements to objects with target and distractor features prior to the choice even when choice accuracy plateaued and learning completed (**Fig. 3C-E**). This extended sampling of feature information supports the notion that exploratory fixational sampling of objects is guided by a representation that is separable from representations guiding exploitative fixations used to choose an object.

Based on the these results we predicted that selection history biases of explorative fixational sampling should be stronger linked with internal states that reflect uncertainty about the relevance of object features when compared with exploitative fixations. We tested this conjecture with a Parallel Belief States model that determines for each object in each trial whether its features are more likely explored, exploited or avoided. This approach confirmed that exploratory fixations of features of objects can be linked to exploratory states of the Parallel Belief State model (**Fig. 4D**), and that fixational sampling during these exploratory states did not show reward history biases (**Fig. 5B,D**). In contrast, exploitative fixations showed prominent and persistent reward history biases of objects even when the chosen objects were in exploratory states (**Fig. 5C**). The model-based analysis confirmed that hidden states about the certainty and uncertainty of the relevance of object features account for the behavioral effects. A parsimonious explanation of this result pattern is that these latent states reflect separable attentional priorities for guiding gaze to visual objects.

### Persistent choices of previous targets reflect habit-like prioritization of previous targets

We found a prolonged bias of subjects to choose objects that contained previous target features that were no longer rewarded. This finding is consistent with a large literature in humans documenting that attention is captured by stimuli that were previous targets (for reviews: (Rusz et al., 2020; Anderson et al., 2021)). Prominent models of attention suggest that selection history controls attentional allocation similar to the control based on top-down goals or bottom-up saliency (Awh et al., 2012; Anderson et al., 2021). This bias of choosing non-rewarded distractor objects is consistently reported to prevail over extensive periods of time, spanning hundreds of trials after which subjects continue to respond faster or more likely to non-rewarded distractors that were targets at an earlier time even when subjects meanwhile have learned novel target associations (Della Libera and Chelazzi, 2009; Liao and Anderson, 2020b, a; Anderson et al., 2021). In computational models, the involuntary choices of abstract features that were previous targets correspond to an involuntary stickiness in the attentional representation that translates into behavioral perseveration independent of ongoing value learning (Balcarras et al., 2016; Chakroun et al., 2020; Tuzsus et al., 2024). This conceptualization is consistent with our result that previous targets not only slowed down the learning process but influenced choices even after learning completed akin to the detrimental selection history effects in visual search studies where choosing non-rewarded distractors is non-adaptive and impairs performance (Kyllingsbaek et al., 2001; Kyllingsbaek et al., 2014; Miranda and Palmer, 2014; Lin et al., 2016; Sha and Jiang, 2016; Qu et al., 2017).

Despite these negative behavioral consequences it has been argued that persistent selection history biases reflect a net adaptive evolutionary mechanism that can improve exploitation of known reward sources in more stable environments by reducing time- and energy-demanding foraging of stimuli by favoring already experienced stimuli (Hills et al., 2015; Anderson et al., 2021). This view resonates with our results and suggests that fixational choices of previous targets reflect a habit-like attempt of exploiting previous goal-relevant, reward predictive stimuli. Consequently, gaze toward previous targets likely indexes a goal-driven attentional priority that is exploiting assumed reward predictions. This interpretation is consistent with our finding that exploitative gaze patterns are prominently influenced by selection history, while exploratory gaze is rather unaffected by selection history (**Fig. 4D**).

### Exploratory gaze reflects attentional prioritization of uncertain objects

Gaze towards stimuli that were previous targets but have become non-rewarded distractors is different from gaze that samples information from objects because they have either uncertain reward values or uncertain feature information. In our task paradigm, subjects fixated one, two or three objects prior to committing choosing one object. These pre-choice fixations were only affected by reward history in the first few trials in a learning block and continued sampling targets and distractors even after objects with the target feature were consistently chosen, i.e. when choice accuracies stabilized. Since these excess fixations occurred prominently on error trials that ended with choosing a non-rewarded distractor (**Fig. 3E**), they likely indicate subjects’ uncertainty about the value or feature of the fixated objects, allowing them to inform the choice process (Konovalov and Krajbich, 2016). Interpretating these pre-choice fixations as being guided by uncertainty highlights that they reflect exploratory fixational sampling of objects. In contrast to the exploitative fixations, exploratory fixations originated in estimates about uncertainty. Uncertainty has been shown to determine gaze shifts to different objects (Ludwig and Evens, 2017). Visual search studies have shown that a stimulus that is informative, i.e. that can reduce uncertainty, attracts gaze even if the gathered information is unrelated to reward outcomes (Ghazizadeh et al., 2016; Pearson et al., 2024; Chow et al., 2025). In these studies subjects were presented multiple stimuli and had to search and choose a target, which involves similar search processes as in our task paradigm. Despite this search goal, fixations were directed to non-instrumental objects in the human studies as they were in our study with nonhuman primates. This similarity suggest that exploratory fixational sampling in our study is reminiscent of so-called uncertainty-modulated attentional capture (Chow et al., 2025).

In summary, the differential effects of selection history on exploratory and exploitative fixations, and the prolonged fixational sampling during exploratory states quantifies evidence for separate attentional priorities acting in parallel during feature-based attention within a multidimensional environment.

## Acknowledgments

This work was supported by the National Institute of Mental Health (R01MH123687). The funders had no role in study design, data collection and analysis, the decision to publish, or the preparation of this manuscript.

## Data and code accessibility

Data and custom programming code for analysis is available upon request.

## Financial Disclosures

The authors declare no competing financial interests.

## Methods

### Ethic Statement

All animal and experimental procedures complied with the National Institutes of Health Guide for the Care and Use of Laboratory Animals and the Society for Neuroscience Guidelines and Policies and were approved by the Vanderbilt University Institutional Animal Care and Use Committee.

### Experimental Procedure and Design

Two male rhesus macaques (Monkey W: 14 yrs / 12.2 kg; Monkey I: 15 yrs / 12.1 kg) performed the experimental task in a sound attenuated chamber. Subjects performed the task with their head position fixed, facing a 21’’ LCD screen at a distance of 63 cm from their eyes to the screen center. The visual display, task, reward delivery, and eye movement responses of the subjects were controlled by the unity-based software platform USE (Watson et al., 2019b). The task required monkeys to choose on every trial one of three objects to earn reward-tokens that were regularly cashed out for fluid rewards. On each trial only one of the objects contained a visual feature that was associated with reward (**Fig. 1A**). Visual objects were 3D-rendered Quaddle objects (Watson et al., 2019a) that varied in blocks of trials in features of two feature dimensions (randomly chosen for each block among the dimensions: body shape, arm style, color, and body surface patterns) (**Suppl. Fig. 1**). A block consisted of 30-55 trials in which one object feature was the designated target feature. Monkeys had to learn which of the six different features that distinguished the three objects was the rewarded target feature through trial-and-error. When they made a choice, a visual halo appeared behind the chosen object in yellow if the choice was correct, and in blue otherwise. Correct choices were followed by green circles (tokens) shown above the chosen object. Tokens moved up towards a token bar and populated previously empty token-slots (**Fig. 1A**). After an incorrect choice and blue halo, grey tokens appeared above the chosen object, moved upwards to the token bar and the grey tokens were subtracted from the already existing green tokens at the token bar. The token bar started at the beginning of the experiment and in each new block of trials (*see* below) with three already available tokens and 5 empty token slots. When monkeys collected eight tokens the token bar flashed, the monkey received water reward, and the token bar was reset to contain three default tokens.

### Block transition conditions

Each experimental session consisted of 36 blocks that lasted 30-55 trials. In each block one feature was designated a rewarded target feature (**Fig. 1B**). The experiment pseudo-randomly determined for the new block whether the newly reward target feature was a feature that was part of the features used in the previous block (‘*Same’* block transitions), or whether the features were new and not shown before (‘*New’* block transitions). The target feature also differed depending on whether it was a feature of the same visual feature dimension as the previous target such as a switch from a red to a green color, or from an oblong to a pyramidal shaped objects (an ‘*intra-dimensional ID*’ block transitions), or whether the target feature was from a different visual feature dimension as the previous target feature, such as a switch from a red color to an oblong shape (‘*extra-dimensional ED*’ block transition) (**Suppl. Fig. 1**). In addition to the Same / New block conditions and the ID / ED conditions each block had one of four different motivational contexts: Within a block, monkeys either earned 2 or 3 tokens for a correct choice, and they either lost 1 or 3 tokens for an incorrect choice, which combined to four distinct motivational contexts (Gain 3/Loss 1; Gain 3/Loss 3; Gain 2/Loss 1; Gain 2/Loss 3). The motivational contexts remained constant within a block and varied pseudo randomly within an experimental session so that each context was equally often assigned to each of the four block transition types (Same/New and ID/ED transition types). The behavioral performance showed that learning speed, i.e. the trials needed to reach criterion performance, varied consistently between the Same/New blocks (**Fig. 1C-F**), and less consistently with the ID/ED, Gain 3/Gain2, and Loss 3/Loss1 block conditions (**Suppl. Fig. 2**).

### Monitoring and analysis of gaze

Gaze was collected at 300 Hz sampling rate with a Tobii TX300 infrared eyetracker. The monitor was placed above the tracker in its standard physical configuration. The active display area was 50.8 x 20.4 cm, with a resolution of 1920×1080 pixels and a 60 Hz refresh rate. Gaze was calibrated with 9-points at the start of each experimental session. Postprocessing classified gaze into five types: saccades, post-saccadic oscillations (PSOs), fixations, smooth pursuits, and undefined using procedures detailed elsewhere (Watson et al., 2019b; Voloh et al., 2020). Algorithms applied for gaze classification first removed artifacts including blinks and off-screen gaze locations from the raw gaze data from each eye individually. Location estimates were averaged between the two eyes. Minor missing data (<= 2 sample or 6.6 ms) was estimated to remain natural patterns using shape-preserving cubic spline interpolation. Then a Savitzky-Golay filter was applied to 20-millisecond windows to smooth the x and y position data without distorting important movement features. After the steps the gaze classification algorithm operates in three stages. First, it detects saccades by identifying sharp increases in angular acceleration. This process begins with calculating acceleration using a smoothing differentiation filter, then establishing adaptive thresholds using median and median absolute deviation to avoid outlier bias. Putative saccadic onsets are defined as points of maximum acceleration crossing the threshold, with offsets at minimum acceleration points. Short saccades (<10ms) are ignored and closely spaced ones are combined. Precise saccade boundaries are refined based on direction deviation and inconsistent directional changes. In the second stage, post-saccadic oscillations (PSOs) are identified following each saccade. Finally, the remaining intervals are classified as either fixations or smooth pursuits. Segments are labeled as smooth pursuits if they meet four criteria: sample dispersion <0.45, consistent direction >0.5, path displacement >0.3, and spatial range >1.5°; otherwise, the segments are classified as fixations.

### Detecting the fixations onto objects and token bar

Preprocessed eye-tracking data were analyzed to identify fixations on objects and the token bar during the time interval between stimulus onset and selection fixation (**Fig. 3A,B**). For each recorded fixation, we calculated the Euclidean distances between the fixation coordinates and the center coordinates of all objects and the token bar. A fixation was classified as being on an object when its coordinates fell within a 120-pixel radius of the object’s center. Similarly, when a fixation fell in the token bar area on the top of the screen is classified as a token bar fixation. To eliminate spurious detections, temporal filters were implemented: object fixations shorter than 50 ms (approximately 1% of the total) were excluded from analysis. Additionally, we merged consecutive fixations directed at the same object when separated by gaps shorter than 20 ms, treating them as a single fixation event.

### Classification of fixation patterns

We classified trials by the order of their valid fixations towards targets and distractors (**Fig. 3A,B**). In total there were 53,468 valid trials, 73.1% of trials ended with a fixation to the target (Tc), and 26.9% of trials ended with a fixation to a distractor (Dc). In total, 97.48% of trials contained fewer than or equal to 4 fixations (3 fixations prior to the final fixation that reflected committing to the fixated target, Tc, or distractor, Dc), 87.38% of trials contained fewer than or equal to 3 fixations, and 58.59% of trials contained fewer than or equal to 2 fixations. Trials were then grouped by their last two fixations (**Fig.’s 3,4,5**) or last three fixations (**Suppl. Fig. S3**), with conditions fewer than 3% of total trials grouped together into “Others”.

### Comparing learning speed for choice accuracy and fixational sampling sequences

We defined the speed of learning in a block as the number of trials needed to reach plateau performance. We evaluated the beginning of plateau performance as 90% of the maximum saturation point of a half-sigmoid fit applied to the accuracy curves (**Fig 1E**). Session-wise saturation points were plotted to compare the learning speed difference of overall choice accuracy and choice accuracy for trials based on the end fixation (**Fig. 3D**). Welch’s t-tests were applied to compare each end fixation pattern to accuracy.

### Parallel Belief State (PBS) Model of Attention

We used the recently proposed Parallel Belief State (PBS) model of attention (Tiesinga and Womelsdorf, 2025) to objectively quanify which object features are considered by subjects for an exploratory or an exploitative eye movement and choice. The PBS model is an input-dependent Hidden Markov Model (Bengio, 1994; Ashwood et al., 2022; Murphy, 2023) to infer hidden states *z*, which reflect the belief about a feature being the target, and determines the state transition during the current trial as a function of an input which represents the choice and outcome on the preceding trial. The mechanics of the model are described in (Tiesinga and Womelsdorf, 2025) and summarized in **Figure 4A**. In brief, the model accounts for observations of whether a feature was present or not in a chosen object by maximizing the probability for the entire set of observed choices (*c*_1:*T*_) across trial 1 to T, and the corresponding hidden states (*z*_1:*T*_) given the inputs (*h*_1:*T*_) and a particular model m with parameters *θ*_*m*_, which is written as: *p*(*c*_1:*T*_, *z*_1:*T*_|*h*_1:*T*_, *θ*_*m*_). The notation *θ*_*m*_ is shorthand for a collection of parameters including the transition and emission matrices as well as the distribution of initial states, which are detailed below. The choice *c*_*t*_ (value 1 if the feature is chosen, that means it was present in the chosen object, and likewise 2 if it is not chosen) on trial t is predicted based on the current input *h*_*t*_ = *c*_*t*−1_ + 2(1 − *r*_*t*−1_) which reflects the choice and outcome (*r*_*t*−1_ = 1, when rewarded and is zero otherwise) on the preceding trial encoded as a number between 1 and 4. This input directly influences the predicted choice via the input-dependent emission matrix 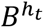 (size: number of choices (2) times number of states n_s_), which links the current hidden state *z*_*t*_ which reflects the longer-term history to a probability of the choice *c*_*t*_ via the matrix element 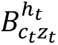 of this matrix. The input dependent transition matrix 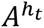 (size n_s_ times n_s_), updates the hidden state to reflect the most recent available choice/outcome, which reflects the learning dynamics of the subject. This new hidden state is not directly accessible for the observer, instead it must be inferred, which yields a probability *γ*_*t*_(*j*) for *z*_*t*_ to be in state j when taking into account both the past and the future.

To optimize the prediction success of the model we maximize the likelihood of the actual choice via an EM procedure. We used the log likelihood (LL) of the entire sequence of choices, normalized by the number of choices, its value should exceed that of chance 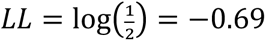 (note that we use a natural e-based log). For each of these data sets we ran the procedure 10 times for different initial values for the parameters (the matrices A, B as well as the prior on the distribution of the hidden states, *π*), to avoid the EM procedure gets stuck in local maxima and performed a minimum of 300 iterations for our analysis.

### Preprocessing the dataset for modeling

We fit the model to choices of the subjects in trials across multiple blocks. In each block one object feature was the target feature and present in one of the three presented objects. Each of the three objects shown per trial had features of two dimensions that distinguished it from the other objects.

One of the features was the designated rewarded target feature across all trials in the given block. To preprocess the data for the model optimization we translated the chosen object and outcome of each trial, for each considered feature, into *c*_*t*_ = 1 when the chosen object contained the feature, and *c*_*t*_ = 2 when it did not. We performed two types of fits, one focused on how a single target feature is learned, the second on how feature belief states change in parallel, whose results are reported in the main text.

For the first fit, we consider each feature separately and find all the blocks in which it was the target, and for each target block include the preceding block, which are then all concatenated. We then concatenate these single feature data sets across all features. The primary reason for doing this is that the feature under consideration has an equal likelihood to be a target or not, so that the model prediction has to be better than a random choice with probability half for each option (hence corresponding to a log-likelihood 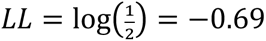). Furthermore, the “exploit” (or “persist”) state, indicating that the feature is chosen persistently because it is the target feature, should occur in a reasonable fraction of trials so that its properties can be well estimated.

In each block there were six features present in the objects, when in a subsequent block one or two new dimensions are introduced (in the condition ‘New’), there are either three or six new features added (see **Suppl. Fig. S1**). For the second fit, we organized the data so that we can track these block transitions, but avoid having to track the states of too many features simultaneously because we will not get appropriate statistics on each state, hence we cut the trials into groups of consecutive blocks in which no more than 12 unique features are present. This means that the set of features (and their index) will vary across these groups. The choices for each of the (at most 12) different features are combined in one big set and fitted by a common transition matrix. This way of combining choices means that the “exploit” state will occur less often for each feature, because it is the target in a small fraction of blocks. This makes fitting additional states reflecting the learning process of target challenging.

We set out to find the best set of parameters for the model, in terms of the dimensionality of the hidden state, *n*_*s*_, the depth of the history *h*_*t*_ (how many previous trials are incorporated, in our standard formulation this was 1) and the best model type (the input dependence of transition and emission matrices). We were guided by our previous analyses (Tiesinga and Womelsdorf, 2025), hence we only explored *n*_*s*_ = 3 and 4 and considered only the case in which the emission matrix was not input dependent. In the first fit there was a difference in number of feature choices (M1: 71074, M2: 28835), which lead for M1 four states to have a lower (better) negative log likelihood (NLL) and BIC than for three states (NLL=0.4131 vs 0.4189, BIC= 59336 vs 59866, for three states there were 29 and for four states there were 55 parameters). However, for M2 the NLL was still lower for four states, but the BIC was higher (NLL=0.3914 vs 0.3958, BIC= 23139 vs 23121), indicating that improvement in fit was not sufficient to compensate for the increased number of parameters.

For the second fit, we again compared 3 and 4 states, but fixed the emision matrix, based on a prior experience (Tiesinga and Womelsdorf, 2025), hence the number of parameters reduced to 26 and 51, respectively. The number of feature choices available for M1 and M2 was 425016 and 212256, respectively. For this case, four states led to a lower NLL and BIC for both subjects (M1: NLL=0.2860 vs 0.2886, BIC=243864 vs 245697; M2: NLL= 0.2747 vs 0.2784, BIC= 117281 vs 118560). However, when we inspected the fraction of trials the feature was chosen in each state, it turns out the “exploit” state was artificially split into two states, with approximately the same fraction of choices. We therefore settled on performing the analysis with three states. We define the states as “*Avoid*”: do not choose the feature (lowest probability the feature is chosen); “*Explore*”: choose the feature with probability 0.33; and “*Exploit*”: (almost) always choose the feature. Each state has therefore a clear interpretation in terms of predicted feature choice.

The parallel belief states comprise the combined states of each of 12 features. We assign a label to each PBS of each trial (**Fig. 4E,F**) based on the distribution of states across features as follows:

- **Exploit target**: target feature is in “persist” or “exploit” state
- **Explore target**: target feature is “explore” state, no others features are in the “explore” state
- **Explore distractor**, one or more non-target features states are in an “explore” state, which can include the target feature
- **Perseverance**, feature corresponding to target of preceding block is in “persist” or “exploit” state
- **Persist in distractor feature**, a different feature (not current target, not previous target) is in an “exploit” state
- **Other**: none of the above

The *Other* state category occurred too rarely to meaningfully separate it further:

## Notes

### Competing Interest Statement

The authors have declared no competing interest.

