## Supplementary Figures for "Two Distinct Attentional Priorities Guide Exploratory and Exploitative Gaze in Parallel"

### Supplementary Figures S1-S4

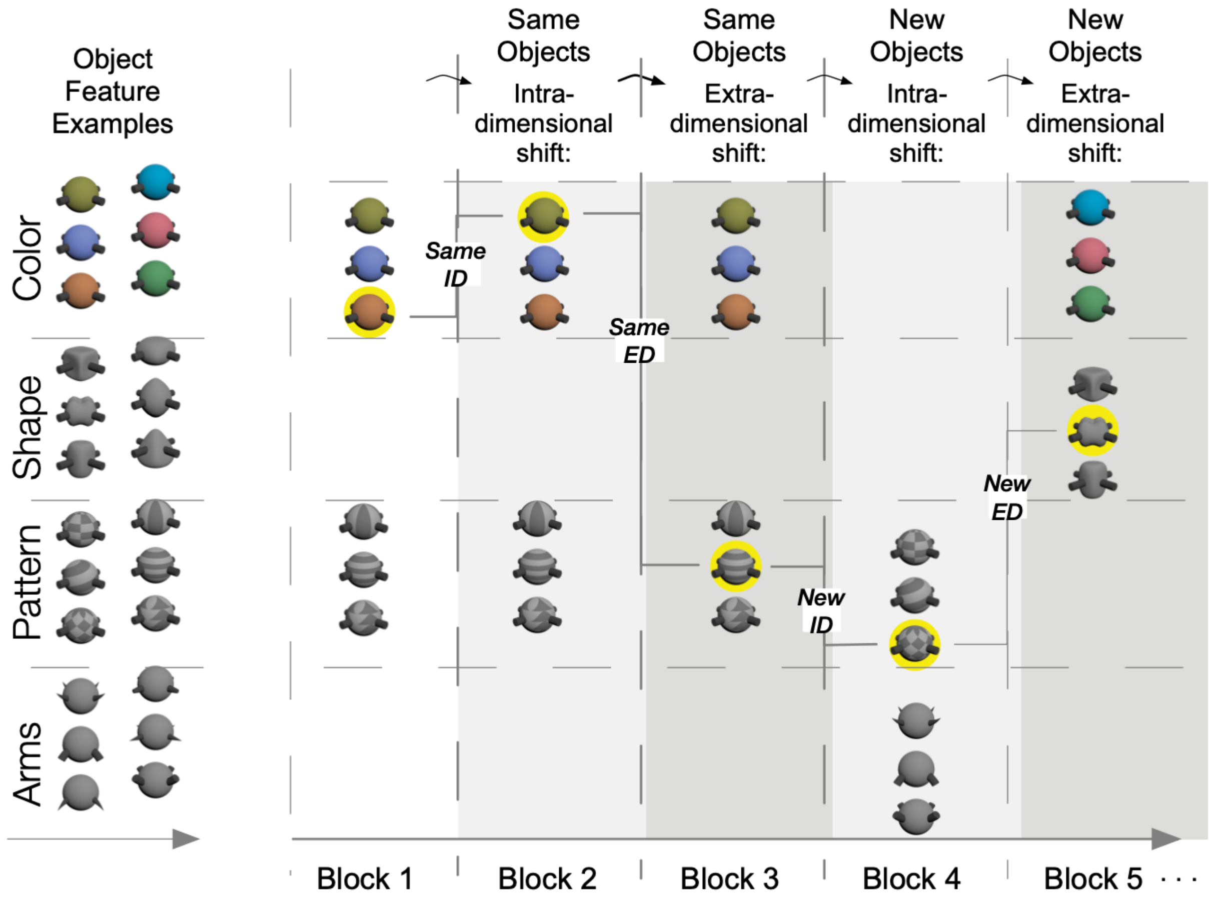


**Supplementary Figure S1.** Illustration of block transitions. Example transitions when the same objects were used and the target feature shifted intra-dimensionally from Block 1 to Block 2 (ID-Same), or when new objects were used and the target feature shifted extra-dimensional from Block 4 to Block 5 (ED-New). Other block transitions show ID-New and ED-Same.

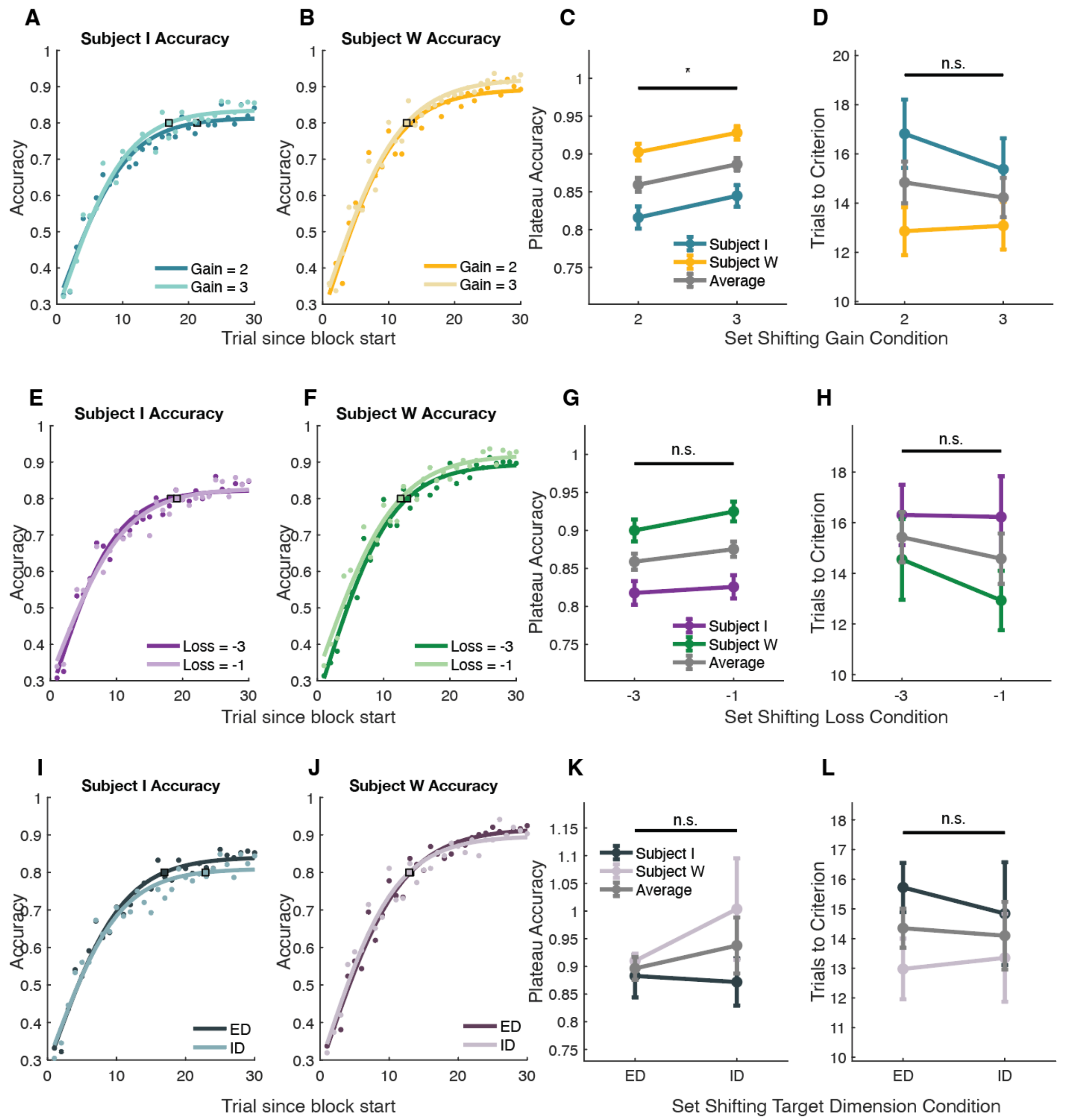


**Supplementary Figure S2. Performance Curves in Blocks with variable gains, losses, and intra-/extra-dimensional target feature transitions.** (**A**,**B**) Proportion of correct choices for subject I (*A*) and W (*B*) increased over trials at similar speed when correct choices led to a token gain of 2 or 3 tokens. Open squares denote the trial at which 90% of maximal plateau performance was reached which was the criterion defining learning. (**C**) Plateau performance was higher when subjects anticipated 3 versus 2 token gains for correct responses. Star denote significance at $p<0.05$. (**D**) Learning speed was similar across conditions. (**E-H**) Same format as *A-D* for blocks in which subjects lost 3 or 1 token for incorrect choices. Plateau performance and learning was on average similar across conditions. (**I-L**) Same format as *A-D* and *E-H* for blocks in which the target feature was from the same dimension as in the previous block (intra-dimensional, ID) or from a different dimension as in the previous block (extradimensional, ED). Plateau performance and learning was on average similar across conditions.

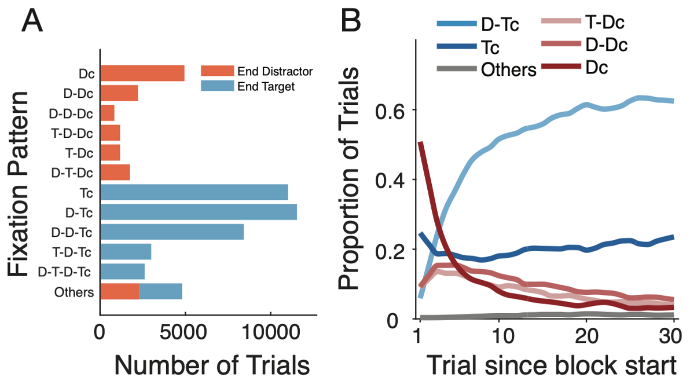


**Supplementary Figure S3. Patterns of fixational sampling and choice for different long fixation sequences.** (**A**) Total number of trials (*x-axis*) with fixation to targets (T) and distractors (D) and fixational choices of the target (Tc) and the distractor (Dc). (**B**) Same data as shown in Fig. 3D: The proportion of fixational sampling sequences (y-axis) across trials in a block for trials with no fixational sampling prior to a choice (Tc, Dc), and for all other trials but considering only the last fixation prior to the choice. (**C**) Same format as B but including only trials in which subjects looked at two objects prior to choosing an object.

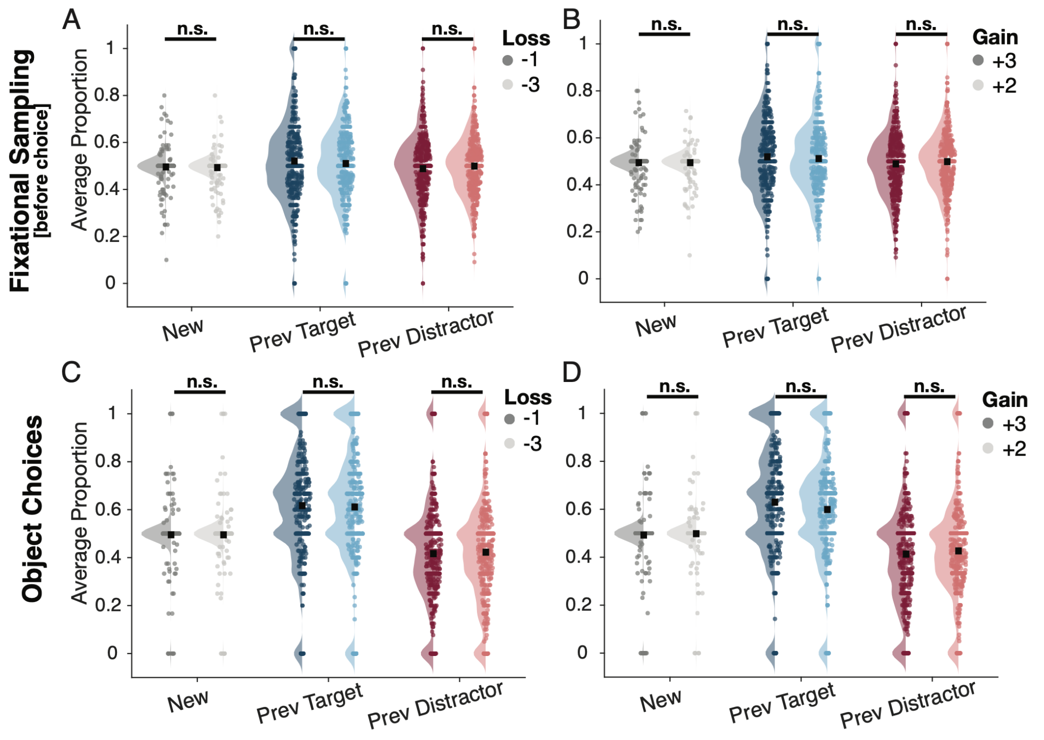


**Supplementary Figure S4. No differences of selection-history effects on fixational sampling and choices in the Gain and Loss conditions**. (**A**) Average proportion of trials (±SE) in which subjects erroneously chose a distractor object with features that were *New* (that were not used in the previous block), or were *Previous Target* features or *Previous Distractor* features in the last bock for blocks with a Loss of -1 (dark color) or -3 (lighter color) for incorrect responses. The two loss conditions showed similar effects (not significant two-sample t-test with a Bonferroni correction). (**B**) Same format as *A* for block conditions with a Gain of +3 (*darker*) and +2 (lighter) for correct responses. (**C-D**) Same format as *A-B* for choices of objects. The motivational conditions showed similar selection history effects.
